# Eco-evolutionary dynamics lead to functionally robust and redundant communities

**DOI:** 10.1101/2021.04.02.438173

**Authors:** Lorenzo Fant, Iuri Macocco, Jacopo Grilli

## Abstract

Microbial communities are functionally stable and taxonomically variable: species abundances fluctuate over time and space, while the functional composition is robust and reproducible. These observations imply functional redundancy: the same function is performed by many species, so that one may assemble communities with different species but the same functional composition. The clarity of this observation does not parallel with a theoretical understanding of its origin. Here we study the eco-evolutionary dynamics of communities interacting through competition and cross-feeding. We show that the eco-evolutionary trajectories rapidly converge to a “functional attractor”, characterized by a functional composition uniquely determined by environmental conditions. The taxonomic composition instead follows non-reproducible dynamics, constrained by the conservation of the functional composition. Our framework provides a deep theoretical foundation to the empirical observations of functional robustness and redundancy.

## Introduction

The staggering taxonomic diversity of microbial communities parallels with their remarkable functional robustness [1, 2]. At the species and strain level, their taxonomic composition is highly variable across communities with similar environmental conditions and over time. This variability is also observed in microcosmos experiments, under very controlled conditions [3]. On the other hand, the functional composition of communities, estimated for instance using metagenomic data [2, 4], appears to be highly reproducible and stable over time. This — only apparent — contradiction strongly suggests that microbial taxa are highly functionally redundant: since many species can perform the same functions, there exist multiple species combinations corresponding to the same functional profile.

While the replicability of the functional community composition is robust and observed across ecosystems, including laboratory experiments with controlled conditions [3], its origin is unknown. This lack of understanding is, in part, because our theoretical understanding of ecological models focuses on species composition. Population abundances are the standard degrees of freedom of mathematical models of community dynamics.

Consumer-resource models are the main modeling framework for microbial communities. Their origin goes back to the classic work of MacArthur and Levins [5], which has been extensively studied and discussed in the following decades [6, 7], mostly to describe the coexistence of a handful of species. Recently, these models have been further extended to consider facilitation through cross-feeding [3, 8, 9], where species change resource availability not only by consumption, but also because they release in the environment the waste products of their metabolism. These models qualitatively describe experimental results [3, 10] and have the flexibility to reproduce patterns observed in empirical microbial communities [11].

Once the parameters of the model are set and an initial pool of species is chosen, populations converge for large times to an equilibrium point. Under some mild conditions, identified over decades of theoretical work [12, 13], consumer-resource models are characterized by a globally stable equilibrium: the steady state is independent of the initial population abundances and resource concentrations. The competitive exclusion principle — one of the most fundamental results of theoretical ecology — limits the number of species that can coexist in a stable equilibrium: diversity cannot exceed the number of resources. While this bound is hard, it is often not realized, as only fewer species can coexist [14, 15].

The number and identity of the species coexisting at equilibrium is in fact determined not only by the ecological dynamics, but also by the initial pool of species. This initial pool of species is often interpreted as the metacommunity diversity: the ecological dynamics unfolds in a local community which is coupled to the metacommunity by rare migrations. Most of the recent progresses in understanding the assembly of large ecological communities have been driven by the assumption of “random” species pools [15–17]. This choice assumes that the parameters characterizing species’ physiological and ecological parameters are independently drawn from some distribution. This assumption implicitly underlies a separation of spatial and temporal scales: the ecological dynamics determining the community composition in the local community occurs independently of the evolutionary processes determining the pool of diversity of the metacommunity.

Instead of assuming a fixed species pool, one can let evolve dynamically individual traits, including the ones specifying their interactions with other individuals and the environment. Classic work in adaptive dynamics [18] has shown how, starting from a clonal population, diversification can evolve under general conditions on frequency-dependent selection. Several works have then studied eco-evolutionary dynamics of interacting populations [19, 20], by allowing individuals’ traits to be subject to mutations and be inherited by the following generation. “Intrinsic” fitness, how fast populations grow in an optimal environment, and niche differences, how the growth of different populations is coupled, both influence community evolution and is their interplay to determine the observed diversity of an evolved community [21].

A key difficulty in interpreting the outcomes of eco-evolutionary dynamics is the fact that there are no natural degrees of freedom to characterize the evolution of the community. The identity of populations, and not only their abundance, is under constant change.

Here we show that the functional composition emerges as the natural variable that characterizes the composition of the community. We consider the broad framework of consumer-resource-crossfeeding models under an explicit eco-evolutionary dynamics, where strains differ in their resources preference and their intrinsic fitness. Higher resource intakes are balanced by a lower efficiency (or equivalently, higher mortality) implemented by a metabolic trade-off [22, 23]. We show that the evolutionary dynamics converge rapidly to a stationary and reproducible functional composition — here defined as the fraction of individuals able to grow on a given resource — which we analytically predict. Interestingly, we show that, once the functional attractor is reached, the strain dynamics is then dominated by fitness differences, implying that functional composition is robust (independent of small fitness differences) and redundant (is obtained under multiple strain compositions).

## Results

The ecological dynamics is defined by standard consumer-resource-crossfeeding equations [8]. In our framework, individuals are characterized by a resource preference vector that determines the intake rate of each of the *R* resources available in the environment (relative to a maximum). An individual with preference *a*_*i*_ = 0 will not consume resource *i*, while an individual with preference *a*_*i*_ = 1 will consume it with a maximum intake rate *ν*_*i*_ [22]. Consumed resources are converted into biomass with finite efficiency (equivalent to an inverse yield). We assume that the yield (or equivalently the death rate, see Materials and Methods) depends linearly on the number of resources consumed: the more resources an individual can grow on, the less efficiently it grows.

Two populations with identical resource preferences can differ in the values of other physiological parameters (e.g., efficiency or mortality). Such differences, which we refer to as intrinsic fitness, determine which population of the two survives when competing. We will use the word ‘strain’ to identify a group of individuals with equal resource preferences and intrinsic fitness.

Resource dynamics are described explicitly. Resources are introduced in the system with a resource-specific rate *h*_*i*_ and consumed by the individuals present in the community. Their concentrations decrease because of consumption but also vary due to cross-feeding. A fraction 1 − *ℓ* of resources consumed by each individual is used for growth, while a fraction *ℓ* is transformed into different resources and released again in the environment [8]. The cross-feeding matrix, with elements *D*_*ij*_, specifies the relative rates of resource transformation (see Materials and Methods for its parameterization). The per-capita growth rate of a strain *μ* is a function *g*_*μ*_(*<u>c</u>*) of the resource concentration *<u>c</u>*, which, in turn, depends dynamically on the population abundance because of consumption and cross-feeding. We consider the following choice

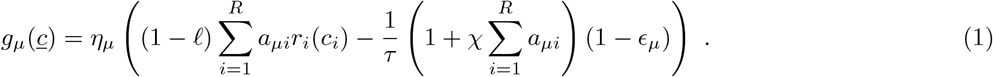

The values of *η*_*μ*_ and *τ* are arbitrary and their choice does not affect the results. The functional form *r*_*i*_(*c*_*i*_) encodes the functional response. Both linear and saturating (Monod-like) functional responses produce the same results (see Fig.S4). The parameter *𝒳* quantifies the fitness cost of consuming one resource and implements the metabolic trade-off. The form of the trade-off generalizes the case considered in [22, 23], which assumes a constant total energy budget devoted to metabolism (i.e. a constant value of Σ_*j*_ *a*_*σj*_). In our case, the total metabolic energy budget is not fixed to a constant but allowed to vary. In the Materials and Methods, we show that the fixed energy budget scenario [22, 23] corresponds to the limit of large values of *𝒳*. Including a constant term in the metabolic cost (set equal to 1 in our framework without loss of generality) is related to a basal cost, related to housekeeping functions. The quantity *ϵ*_*μ*_ determines the intrinsic fitness value.

In our framework, both resource preferences and intrinsic fitness values are subject to mutations and evolution. We consider different implementations of the mutational steps (e.g., including different scenarios for the relative rate of Horizontal Gene Tranfer, see Materials and Methods) which, however, do not affect the results. The timescales between two successive successful mutations is comparable with the ecological timescale, set by the ecological dynamics. We assume that a mutation of the resource preference always corresponds to a mutation of the intrinsic fitness, which is drawn at random from a fixed distribution with width *ϵ*. The parameter *ϵ* sets the typical difference of intrinsic fitness values between two individuals. We focus on the case of small fitness differences, and we extensively explore the effect on the eco-evolutionary trajectories of increasing the value of *ϵ*.

Fig 1 shows a sample of eco-evolutionary trajectories resulting from our framework. Starting from a clonal population, a diverse community is rapidly assembled. Strain abundances change abruptly following successful invasion events and keep changing over the whole duration of the simulations.

**FIG. 1:**
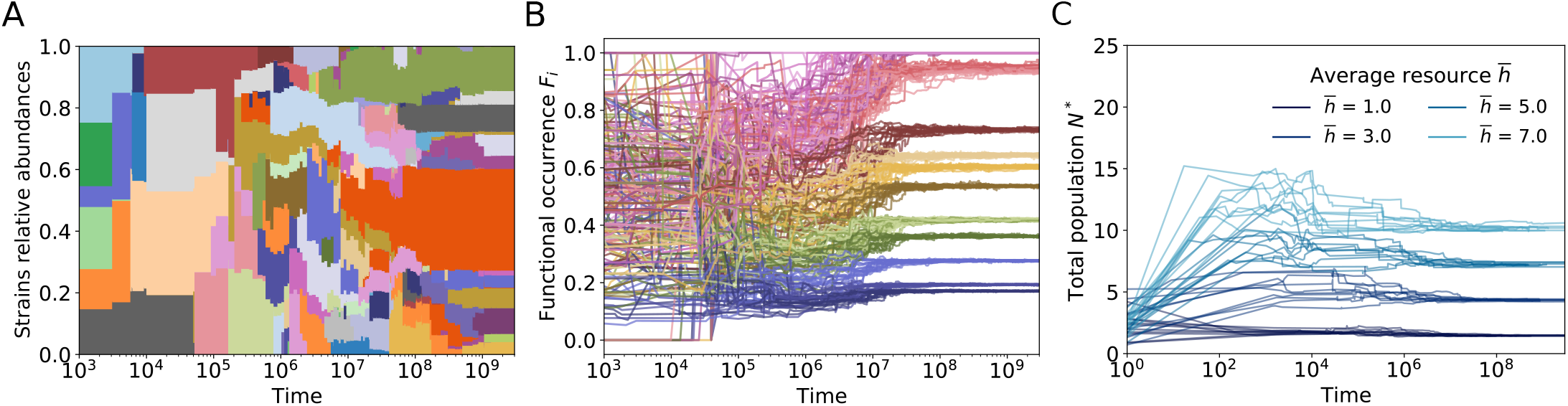
Stability of functional occurrences *F*_*i*_ for communities evolving under a consumer-resource model. The system is initialized with a small number of initial random strains, chosen so that each gene is present at least once. The system evolves in a chemostat with fixed resources input. When equilibrium is reached, one mutant is added to the batch. The chemostat then equilibrates to a new fixed point and the procedure is repeated until function and biomass reach stability. **A**: Time evolution of the strains relative abundances for one realization of the system. **B**: Time evolution of functional occurrences for three different realizations of the system. 15 resources are given. **C**: Time evolution of the total biomass of the system. 20 realizations of the system are shown for each value of average resources income.

The final community structure is remarkably simple if, instead of analyzing strain abundances, we focus on its functional composition. We define functional occurrence *F*_*i*_ as the fraction of individuals able to grow on *i* (i.e., with *a*_*i*_ = 1). After a short transient, the functional occurrences and the total biomass *N* relax to the respective stationary values 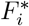 and *N* ^*^, which are very reproducible across different realizations.

Two phases characterize therefore the eco-evolutionary dynamics. The first one is an initial-condition-dependent transient, where the community structure is mainly shaped by rapid invasions. In the second phase, conversely, the community has converged to a stable functional composition, which we will refer to as “functional attractor” in the following, and slowly evolves reaching the final strain-level equilibrium.

The total biomass converges to a constant value during the second phase of the eco-evolutionary dynamics, which affects therefore only the relative abundance of strains. The sequence of invasions and extinctions of strains is determined by the interplay of fitness differences *ϵ*_*μ*_ and niche differences, related to the dissimilarity of the resource preferences. Importantly, the trajectories of strain abundances are effectively restrained to occur on the low(er)-dimensional space determined by the constraint enforced through the functional occurrences 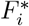. In this second phase the community has thus reached a “functional maturity” and the subsequent evolution only affects strain composition while leaving unaltered the functional one.

The stability and reproducibility of the functional attractor suggest that it is possible to predict analytically its properties. We considered a toy model of the eco-evolutionary dynamics which aims at mimicking the effective exploration of the phenotypic space performed by mutations. In particular, we consider only the ecological dynamics, initialized with an infinitely large species pool, which encompasses all the possible strains (i.e., the 2^*R*^ possible resource preferences). A similar approach has been considered to study a simpler version of the model [22] (corresponding to the limit *𝒳* ≫ 1 and no cross-feeding). The toy model further postulates a timescale separation between resource and population dynamics [22, 23], which is not assumed in the full eco-evolutionary dynamics.

The consumer-resource-crossfeeding model with infinite pool of diversity and no intrinsic fitness differences can be analytically solved. In the Materials and Methods we show that the stationary functional occurrences 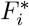 and the total biomass *N* ^*^ are given by

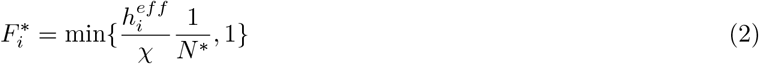

and

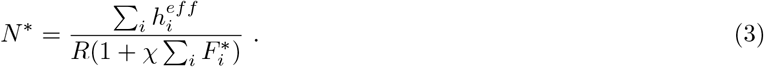

The parameter 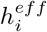 is the effective resource inflow in the system, which is given by the combination of resources that are externally supplied and the ones produced via cross-feeding. This quantity is in simple linear relation to the inflow rate of externally provided resource *h*_*i*_ through the cross-feeding matrix *D* (see Materials and Methods).

The analytical calculations are based on many simplifying assumptions (infinitely large pool of diversity, not explicit resource dynamics, absence of fitness differences) which do not strictly hold for the more complex setting of the eco-evolutionary model. Nevertheless, Figure 2 shows that the predictions of eq. 2 and eq. 3 accurately describe the outcomes of the eco-evolutionary dynamics.

**FIG. 2:**
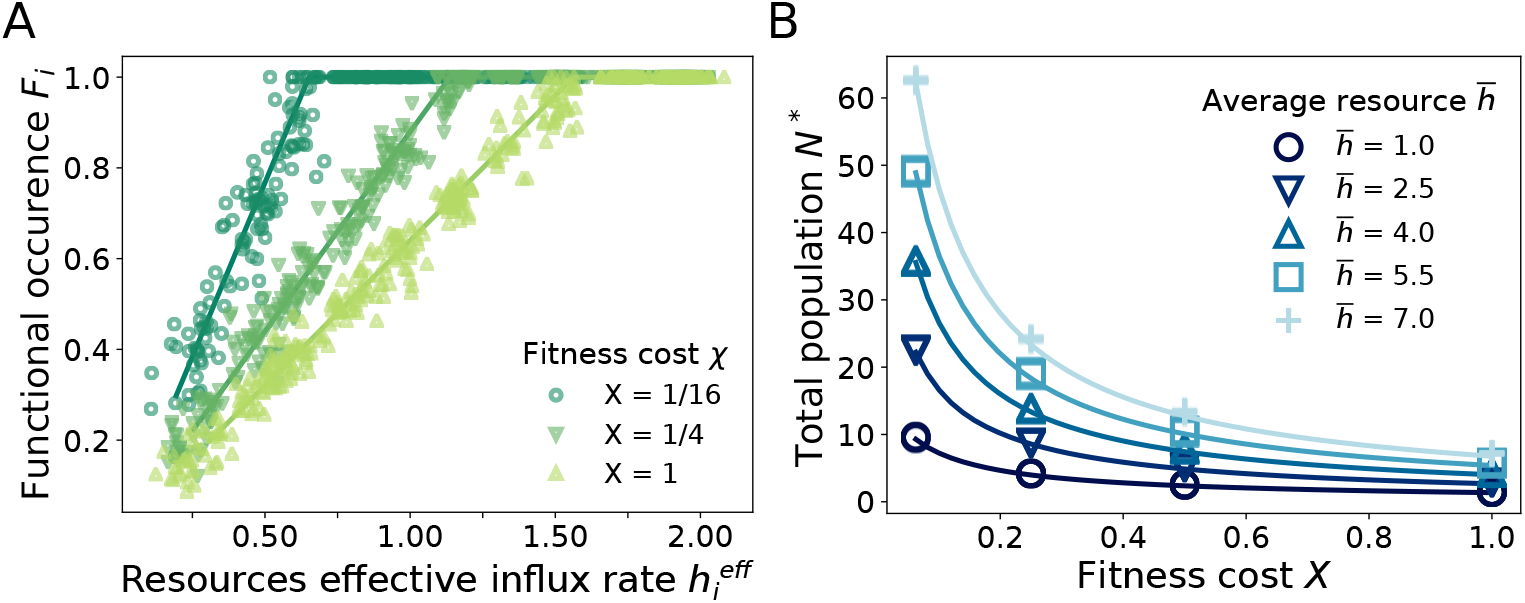
The theoretical predictions given by eq. 13, 14 (solid lines) are reproduced by numerical integration of eq. 4, 5 (markers). **A**: Occurrence of the phenotypes *F*_*i*_ as a function of resource income rates *h*_*i*_. According to equation 13 the most abundant resources (core) are consumed by all strains (*F*_*i*_=1) while the remaining ones only by a fraction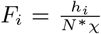. Notice that increasing *𝒳* reflects in a decrease of the number of core resources. **B**: Dependence of the total equilibrium population *N* ^*\**^ (biomass) on the value of *𝒳*. Here we find a dependence on the average value of the resources incomes 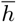, which is absent for quantities in panel **A**. In each figure are represented 20 different noise realizations solutions of the system for each *𝒳* (**A** and each 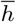 (**B**)

Resources can be partitioned in two groups based on their effective influx rate 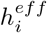. If the influx rate is larger than a critical value 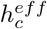, then the ability to metabolize that resource is a “core” function, shared by all the individuals in the community (i.e.,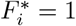). The value of 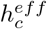 depends on both the spread of the effective influx rate (the variability among the *h*_*i*_) and the metabolic cost *χ*. The higher the metabolic cost and the variability, the higher the critical influx rate threshold 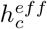 and, consequently, the fewer the core resources.

At equilibrium, the resources with an influx rate below the critical threshold (i.e., the non-’core’ resources) are consumed only by a fraction of the individuals. A linear relation links the functional occurrence 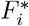 with the effective resource influx rate 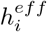. The slope of this relation is simply linked to the metabolic cost and the total biomass, being equal to (*𝒳N* ^*^)^−1^ (see Materials and Methods). Combining equation 2 and 3, one can obtain an explicit expression for *N*^*^. Figure 1B shows that the analytical expression for *N* ^*^ (as a function of the metabolic cost *𝒳*) correctly matches the observations of the eco-evolutionary dynamics.

An emerging feature of the present framework is that the functional composition of communities is extremely robust to fitness differences. We further explore this aspect by considering community response to variation in intrinsic finesses. This variation mimics the temporal or spatial heterogeneity of environmental factors that influence growth, such as abiotic factors (temperature, pH, salinity, etc.) or phages with different host ranges.

We consider two complementary scenarios, which aim at exploring cross-sectional (across communities) and longitudinal (over time) variation. In the former case, we compare the eco-evolutionary outcomes of several communities that share the same resource input but have independent intrinsic finesses. Two individuals with the same resource preference will have uncorrelated intrinsic fitnesses in two different communities. The latter case assumes instead that intrinsic fitness fluctuates over time with a typical autocorrelation timescale (see Materials and Methods). Over time ranges shorter than the autocorrelation timescale, intrinsic fitness is approximately constant. Over times larger than the autocorrelation timescales, the intrinsic fitness decorrelates and becomes an independent variable.

Figure 3 shows the strain and functional composition of communities in the two scenarios described above. The strain composition strongly differs across communities or overtime, being highly sensitive to small intrinsic fitness differences. On the other hand, the functional profile is left largely unaffected by fitness variation. These observations clearly show that functional redundancy naturally emerges in complex consumer-resource-crossfeeding models, closely reproducing the phenomenology observed in microbial communities [2].

**FIG. 3:**
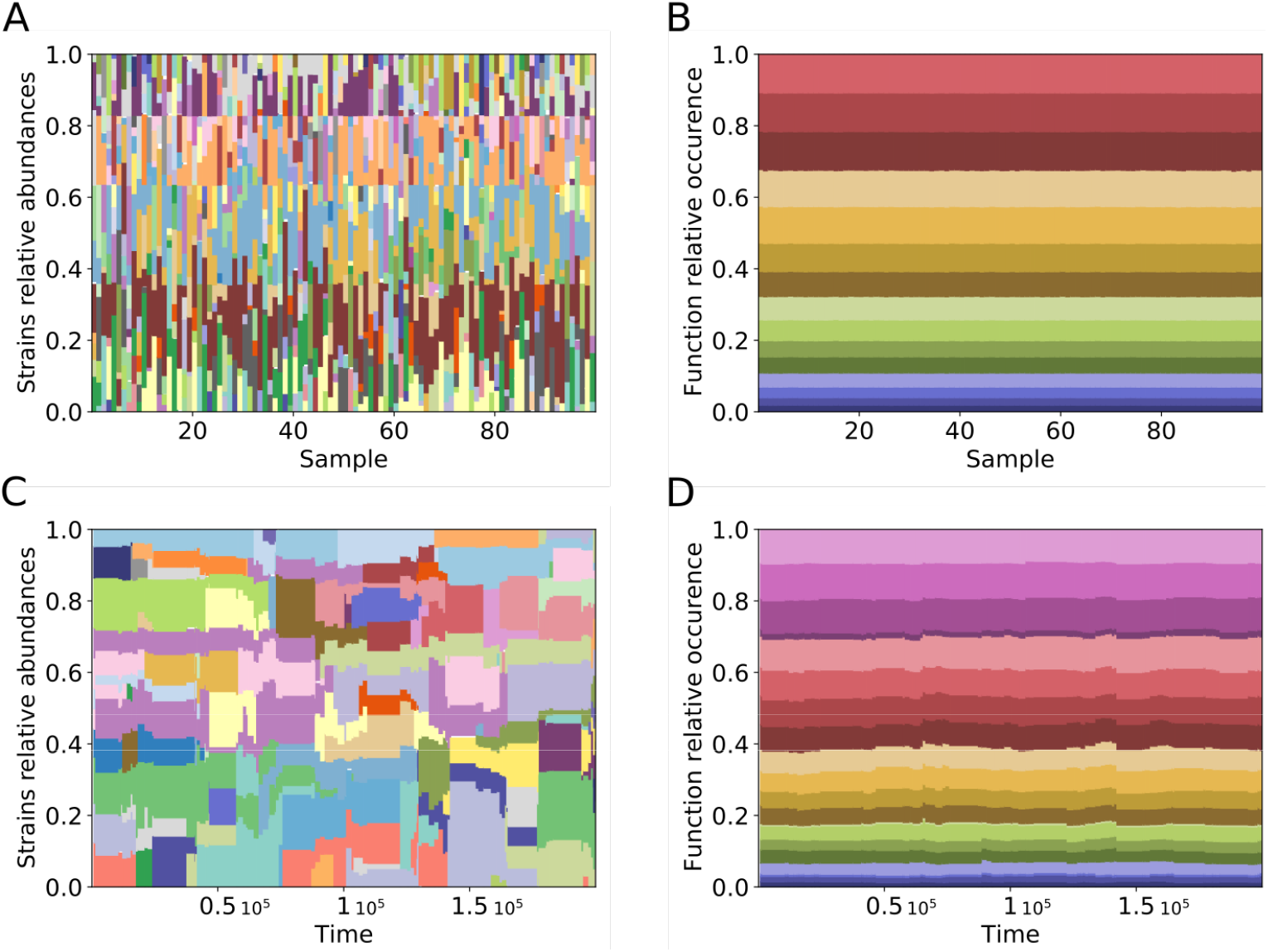
Fitness differences allow to demonstrate how functional stability is encoded within the model. While the populations/taxonomies becomes highly variable and heterogeneous, the functional composition is preserved and unaffected by stochasticity. **A,B** show a collection of equilibrium configurations of systems with different realisations of a static noise. **C,D** show the evolution of the composition of a system with a dynamically varying noise.

## Discussion

Our results shed light on the composition of large ecological communities. When the pool of diversity is not a-priori constrained but is instead allowed to evolve, the complex ecological dynamics can be decomposed in a fast, predictable, phase and a slow one, contingent on the (small, yet relevant) fitness differences. The community composition rapidly converges to a set of solutions, determined by resource availability. The following dynamics are constrained on that subspace of solutions and is governed by the difference in relative fitness. Remarkably, this separation of fast and slow components directly map into functional and taxonomic composition: the former is robust and governed only by effective resource influx rates, the latter is constrained by function, but free to move along functionally equivalent directions.

The functional robustness and functional redundancy are the direct consequences of the existence of the two dynamics phases that directly map onto taxonomic and functional variation. Functional robustness, the observation that functional composition is stable over time and across communities, originates from the existence of a functional attractor of the eco-evolutionary dynamics. While intrinsic fitness differences are small, they are not negligible as they determine the taxonomic composition within the functional attractor. Variation of the intrinsic fitness leads to functional redundancy, high taxonomic variability with conserved functional profile.

The assumption of small intrinsic fitness differences is critical for the observations of functional robustness and redundancy. Increasing the magnitude of fitness differences affects also the functional profile. Typical intrinsic fitness differences of 1% do not alter substantially the functional composition (see Figure S1). Larger differences (of the order of 10%) disrupt the structure of the functional attractor: the functional composition is determined by the resource preferences of the individuals with the largest intrinsic fitness and the functional profile becomes largely decoupled from the resource input. Differences of 0.1% and smaller are indistinguishable from the analytical prediction and the functional profile closely matches the one predicted by the resource influx rates.

The existence of these different regimes, where the functional composition is or is not affected by intrinsic fitness differences, strictly relates to the identification of the limiting factors shaping the communities. As mentioned before, in fact, the *intrinsic* differences could be due to abiotic factors, but also to limiting factors (such as phages) other than resource availability. If resources are limiting, we can expect that other factors have a minimal effect on strains’ success. On the contrary, if resources are not the limiting factors and other mechanisms that determine strains’ growth and decline, the distribution of the functional preferences in the population will not be robust, as it will be subject to the fluctuations of the other limiting factors. Our framework could be extended to explicitly include the factors responsible for intrinsic fitness differences (e.g. phages).

The metabolic trade-off is an essential ingredient of our framework. We implemented it as a fitness cost that is linear in the total rate of resource consumption. This choice generalizes models with fixed total rate [22, 23] by including a basal maintenance cost. This basal cost becomes negligible if the cost per gene, relative to the basal cost, becomes very large. The presence of a non-zero basal cost determines the existence of core resources, whose consumption is shared by all the individuals in the community. The form of the functional attractor is a mathematical consequence of the linearity of the metabolic trade-off. For linear trade-offs, the functional attractor is fully specified by the functional composition and is, in the limit of negligible fitness differences, independent of how functions are distributed across species. Non-linear trade-offs [24] could, in principle, affect the properties of the eco-evolutionary attractor. In the Materials and Methods, we explicitly consider both super-linear and sub-linear trade-offs and show that our results are left qualitatively unvaried. The taxonomic composition is largely affected by fitness differences, while the functional composition is robust. The stable functional composition display core and non-core resources, which are (at least approximately), linearly related to the effective resource influx rates.

A remarkable aspect of our framework is that functional composition — as opposed to taxonomic composition — naturally emerges as the relevant, reproducible degree of freedom suited to characterize ecological communities. Our results demonstrate in fact that the emergence of a stable and reproducible functional composition is a universal feature of consumer-resource-crossfeeding models [3]. This property is likely to hold more generally and not to be restricted to consumer-resource systems or microbial communities. We expect that a similar approach could be developed to study mutualistic communities or pathogen dynamics.

## Materials and Methods

### Definition of the model

We consider a consumer-resource model in presence of cross-feeding [8], which describes the dynamics of population abundances *n*_*σ*_ (for *σ* ∈ 𝒮) and resources concentration *c*_*i*_ (for *i* ∈ *R*). Changes in population abundance are defined by

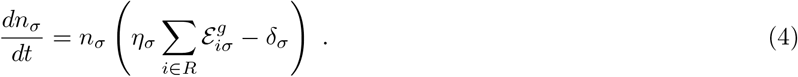

where *δ*_*σ*_ is a death term and *η*_*σ*_ is the efficiency of the conversion of energy into biomass. 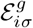 is the energy flux used for strain *σ* to grow from metabolite *i*. We can similarly define the energy flux 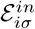 into a cell from resource *i* and the energy released in the environment by the cell 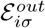 in the form of other metabolites obtained from from metabolites of type *i*. The associated dynamics of resource concentration *c*_*i*_ is defined by

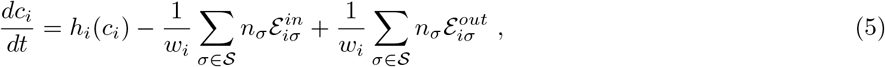

where *w*_*i*_ defines the conversion between energy and concentration of resource *i*. The function *h*_*i*_(*c*_*i*_) specifies the dynamics of resource concentration in absence of consumers.

We assume that energy fluxes used for growth are a constant fraction 1 − *ℓ* of the total ones: 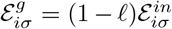. The energy fluxed from secreted metabolites is then given by 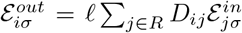. The cross-feeding matrix element *D*_*ij*_ defines energy conversion between resource *j* and resource *i*. Energy conservation implies Σ_*i*_ *D*_*ij*_ = 1.

The energy flux 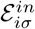 takes the form

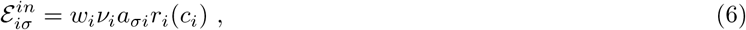

where *r*_*i*_(*c*_*i*_) is a non-decreasing function of the concentration of resource *i* and *ν*_*i*_ is the maximal intake rate of resource *i*. The elements *a*_*σi*_ ∈ [0, 1] measure the intake rate of metabolite *i* strain *σ* relative to the maximum *ν*_*i*_. Here we focus on the case of externally supplied resources *h*_*i*_(*c*_*i*_) = *h*_*i*_, which assumes that dilution is negligible when the total population is around the carrying capacity [23].

We consider a metabolic trade-off by assuming for death rates and yield the expression

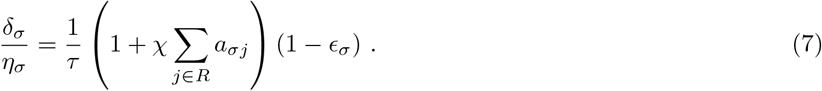

Without loss of generality, we can set the timescale *τ* = 1 equal for all strains, as differences in *τ* can be reabsorbed in the definition of *ϵ*_*σ*_. In the simple setting of *a*_*σj*_ ∈ {0, 1}, the parameter *𝒳* measures the cost of being able to metabolize each metabolite (*𝒳R* is the fitness metabolic cost of a generalist).

### Functional attractor

In the eco-evolutionary simulations, we always consider resources and populations changing over a similar timescale. To make analytical progress we approximate the full dynamics with the effective one obtained by assuming timescale separation — i.e. resource concentrations equilibrate faster than the changes in population abundances. We underline that we assume the separation of timescales only as an approximation, for the purpose of predicting analytically the outcomes of the numerical simulations, which are always obtained with explicit resource dynamics. In this case, one can effectively describe the dynamics of populations as

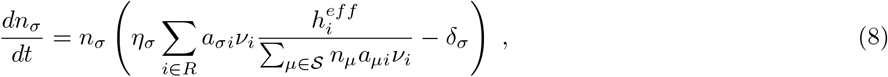

where 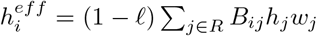 and the matrix *B* = (*I* − *ℓD*)^−1^. It is useful to notice that, in the limit *𝒳* → ∞ and 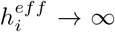 (such that the ration 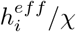 is finite in the limit) reduces to the model with constant total energy budget [22, 23]. It is known [22] that

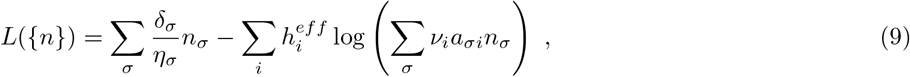

is a Lyapunov function. With our choice for the metabolic trade-off (7), such functional can be conveniently rewritten as

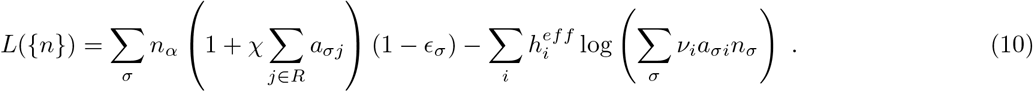

We then introduce the total population size *N* = Σ_*σ*∈𝒮_ *n*_*σ*_ and define the functional abundances *F*_*i*_ as

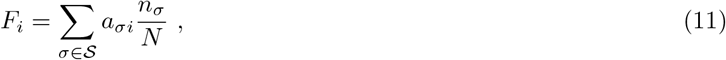

which correspond to the fraction of individuals that are able to metabolize resource *i*. Interestingly, and surprisingly, when *ϵ*_*σ*_ = 0, the Lyapunov function can then be written as function of *N* and {*F*} alone:

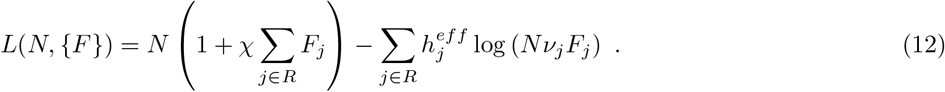

The fact that the Lyapunov function depends only on the total biomass and the functional profile already suggest, even if it does not imply, that functional abundances are the relevant variable for the study of community composition.

By minimizing the function over *F*_*i*_ in [0, 1] one obtains

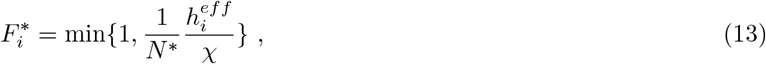

where the total biomass is the solution of

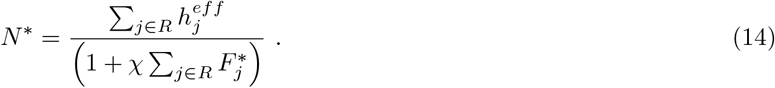

These equation can be solve iteratively, starting from *F*_*i*_ = 1 ∀*i* and 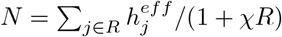.

In the case with no intrinsic fitness differences (*ϵ*_*σ*_ = 0), the equilibrium solutions are identified by equations 13 and 14. For a given system, a fraction of resources will be core resources, i.e. shared by everyone 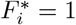. These core resources are the ones for which 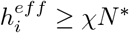.

### Eco-Evolutionary dynamics

The mutation probability of a preference of resource *i* in strain *μ* depends on whether *μ* consumes or not *i*. The rate *U*_−,*i*_ at which a mutant 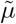 stops consuming resource *i* (the parent has *a*_*i*_ = 1 and the mutant *a*_*i*_ = 0) is constant, independent of *i*, and equal to *U*_−_. The rate at which a mutant starts consuming a resource *i* (the parent has *a*_*i*_ = 0 and the mutant *a*_*i*_ = 1) equals to *U*_+,*i*_ = *U*_+_(*P*_*h*_*F*_*i*_ + *P*_*dn*_). The quantity *P*_*h*_ is the probability that an addition happens because of horizontal gene transfer, while *P*_*dn*_ = 1 − *P*_*h*_ the probability of “de novo” mutations. The rate of horizontal transfer is proportional to the frequency *F*_*i*_ of that allele in the population, while the rate of a de-novo mutation is independent of *i*.

The rate at which the resource preference *i* mutates in strain *μ* is then equal to

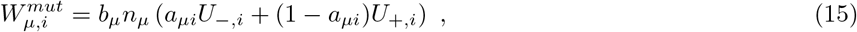

where *b*_*μ*_ is the per-capita birth rate on strain *μ*, which is equal to

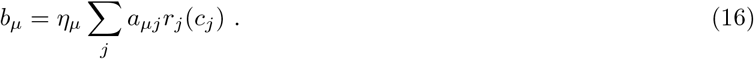

In theory one could expect a new mutant to have abundance 1. The initial phase of its dynamics is then dominated by demographic stochasticity, with many mutants going to extinction despite having a positive (average) growth rate. In our framework, we do not consider this effect of demographic stochasticity explicitly, but we include it effectively. Since the initial abundance of the mutant 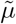 is a small fraction of the total population, its stochastic dynamics can be approximated by a stochastic exponential growth. In this regime, the per-capita birth rate of the mutant is given by 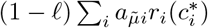, where 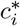 is the concentration of resource *i* prior to the mutant arrival. The per-capita death rate of the mutant reads 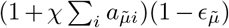. Under the assumption of a stochastic exponential growth the survival probability is given by

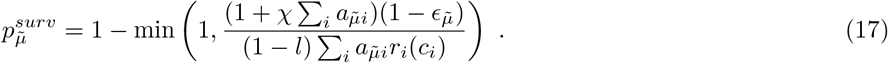

The strain intrinsic fitness values *ϵ*_*σ*_ are independently drawn from a Gaussian distribution with mean 0 and standard deviation *ϵ*.

By calculating all these quantities for all possible mutations of all existing strains, one obtain the rate of invasion 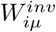 of a mutant 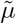 which is obtained by changing the resource preference of strain *μ* for resource *i*. The rate of invasion 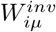 reads

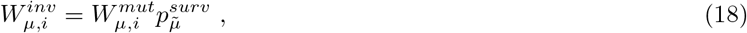

where the mutant 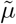 differ from *μ* in the resource preference *i*.

We simulate the eco-evolutionary dynamics as a sequence of discrete small time steps Δ*t*. After a step of integration of equations 4 and 5 we update the values of 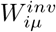, as they depend on strain abundances, and checked whether a mutant appeared. Each mutant, identified by the parent strain *μ* and a resource *i*, has probability 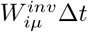 to invade. If such an event occurs, the new mutant is introduced with an initial relative density equal to 10^−5^.

If no mutations appear for a long enough time, the ecological dynamics (obtained by integrating equations 4 and 5) reach an equilibrium point, identified numerically when the absolute value of the population growth rate is lower than 10^−4^. If the strain abundances are not changing, also the rates of invasions 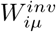 are constant in time (until the next successful invasion), and one can use a Gillespie algorithm. The time of the next successful invasion is drawn from an exponential distribution with average 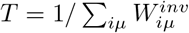. The probability that the new mutant will replace strain *μ* differing in resource preference *i* is simply *TW*_*iµ*_.

### Choice of parameters and sensitivity analysis

The results presented in this paper were achieved using generic parameters, whose details can affect the distribution of taxa or relaxation time but not the macroscopic observables that characterize the functional attractor. In order to quantify the convergence to the functional attractor, we measure the discrepancy between the functional composition of the community during its eco-evolutionary trajectory and the functional composition predicted by equations 13 and 14. As a measure of the discrepancy, we consider the Kullback-Leibler divergence between the normalized functional profiles

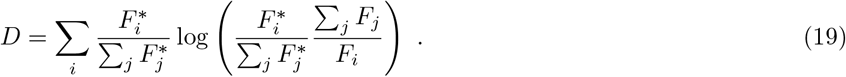

The divergence *D* is equal to zero if and only if the functional composition of the community (quantified by the *F*_*i*_) matches the analytical expectation, i.e. if 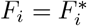 for all the *i*.

We considered *η*_*σ*_ = *ν*_*i*_ = *w*_*i*_ = 1 in eq. 4 and 5. These choices do not affect the results, as they do not affect the ecological fixed point and its stability property (up to a rescaling of the abundances and concentrations). The timescale *τ* was also set to 1, without loss of generality.

In the main text we considered *r*_*i*_(*c*_*i*_) = *c*_*i*_. In the Supplementary Materialswe explore the effect of non-linear intake functions *r*_*i*_(*c*) by considering a Monod-like form *r*_*i*_(*c*_*i*_) = *μ*_*max*_*c*_*i*_*/*(*c*_*i*_ + *K*_*s*_) with different values of *K*_*i*_. Figure S4 shows that the value of *K*_*i*_ has no effect on the functional composition of evolved communities.

The intrinsic fitness on any new mutant was drawn from a Gaussian distribution with mean zero and variance *ϵ* ^2^, independently of the fitness of the parent. In the main text we considered = 0.001. Fig S1 explores the sensitivity of the results to the magnitude of the noise. Much larger values of noise (of the order 0.1) often disrupt the properties of the manifold. For instance, strains not consuming core resources are still able to survive because of high intrinsic fitness. For intrinsic fitness differences with a width of the order of 10^−2^, the functional composition converges to the analytical prediction, which becomes more and more accurate as fitness differences decrease.

The strength of cross-feeding *ℓ* has no effect on convergence to the functional attractor (see Fig. S2). The cross-feeding matrix *D* has been chosen following [11]. Entries were extracted according to a Dirichlet distribution, where resources are in three classes. We considered an effective sparsity of *s* = 0.1. The fraction of resources remaining in the same class was *f*_*s*_ = 0.7 while the ones going to the waste class is *f*_*w*_ = 0.28. The structure of the cross-feeding matrix *D* does not affect the stationary functional composition of the community. Fig. S2 compares a fully random *D* with the ones proposed in ref. [11] and used in the text, observing no difference in the results.

Figure S3 shows that the outcomes of the evolutionary trajectories are independent of frequencies of the different mutation steps. We varied the (average) total mutation rate *U*_*tot*_ = (*U*_+_ + *U*_−_)*/*2, the ratio *U*_−_*/U*_+_ between mutation leading to deletions of resource preferences (with rate *U*_−_) and the ones leading to additions (*U*_+_), and the ratio *P*_*h*_*/P*_*dn*_ between horizontal gene transfer and de-novo mutations. While the total mutation rate, and partially the ratio *U*_−_*/U*_+_, affected the evolutionary trajectories and speed of adaptation, none of these parameters affected the convergence of the functional composition to the predicted attractor.

Both the eco-evolutionary simulations and the analytical approximation are based on the assumption that the metabolic cost is linear in the consumed resources, as expressed in equation 7. In general, one could assume a non-linear tradeoff [24] that takes the form

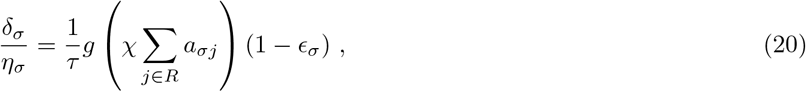

where *g*(*z*) is an arbitrary non-linear, monotonically increasing, function. We considered the outcomes of the evolutionary trajectories in the case of a super-linear cost (*g*(*z*) = (1+*z*)^2^, in Fig. S5) and a sub-linear cost (*g*(*z*) = log(1+*z*), in Fig.S6). In both scenarios, the functional composition converges to reproducible values, minimally affected by fitness differences. On the other hand, the taxonomic composition is much largely affected by fitness differences. Similarly to the linear metabolic cost functions, some resources correspond to core-functions 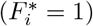 while the functional occurrences 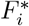 of non-core resources are linearly related to the effective influx rates 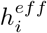. These evolutionary outcomes, obtained under non-linear metabolic costs, confirm the generality of our results beyond the linear metabolic cost case.

## Acknowledgments

We thank M. Tikhonov for insightful discussions and P. Lechon and M. Corigliano for critical reading of the manuscript.

