## Supplementary Figures for "Eco-evolutionary dynamics lead to functionally robust and redundant communities"

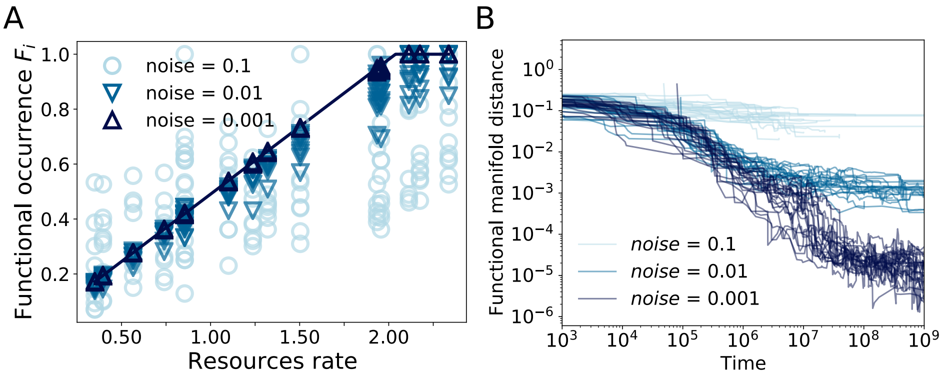


Supplementary Figure S1: Noise amplitude on fitness affects the convergence to the functional manifold. 20 realizations for three different amplitudes, $\epsilon = 10^{-3}$ (dark blue), $\epsilon = 10^{-2}$ (blue) and $\epsilon = 10^{-1}$ (light blue). **A** final functional occurrences of the samples. In the case of $\epsilon = 0.1$ the results falls very far from the noiseless theoretical predictions. **B** distance from the manifold as a function of time. The distance is calculated as $d = -\sum_i \tilde{F}_i^* \ln\left(\frac{\tilde{F}_i}{\tilde{F}_i^*}\right)$, where $\tilde{F}_i := \frac{F_i}{\sum_i F_i}$. In all simulations all the other parameters were set to the same values used in the main text.

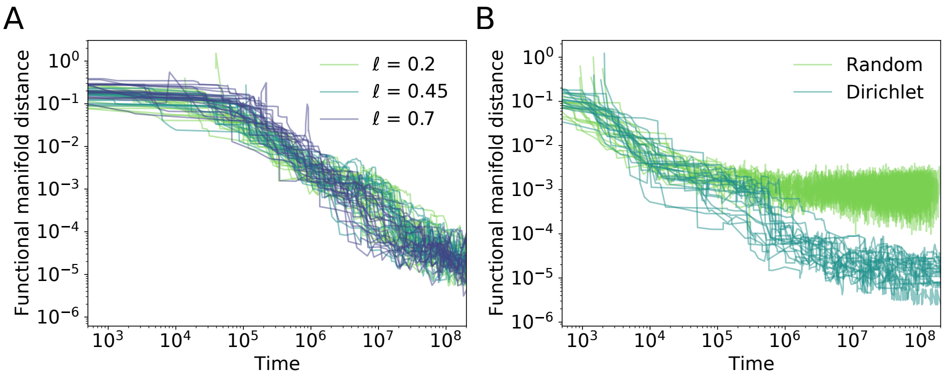


Supplementary Figure S2: Cross feeding effect on the convergence to the functional manifold. The shape and intensity of cross-feeding affect neither the final distance from the manifold nor the path used to reach it. 20 realizations for every choice of the parameters are shown. **A** shows the effects of the amplitude of cross-feeding $\ell$. Three values are here considered, ($\ell = 0.2$) in light-green, $\ell = 0.45$ in green and $\ell = 0.7$ in blue. **B** Difference of convergence behavior in presence of a random cross-feeding matrix and a Dirichlet distributed matrix. The distance is calculated as in Fig. S1 In all simulations all the other parameters were set to the same values used in the main text.

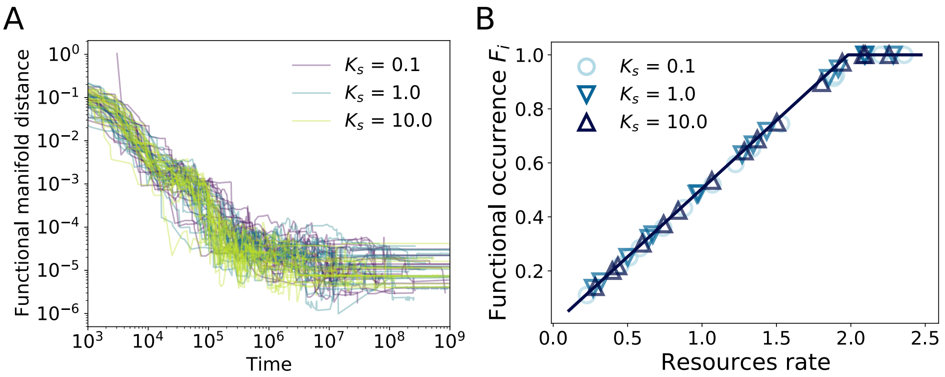


Supplementary Figure S4: The properties of the manifold are insensitive on the choice of the function $r_i(c_i)$ of Eq. (1), i.e. which species consume the resources does not affect the convergence to the functional manifold. In particular both the linear response ($r_i(c_i) = c_i$) and the Monod response ($r_i(c_i) = \mu_{max}\frac{c_i}{K_s+c_i}$) bring to the manifold with the same behavior. In **A** we show the time evolution of the distance from the theoretical manifold for 20 trajectories for every choice of $K_s$ and in **B** the functional occurrence for one realisation for each $K_s$. Such a parameter is not determinant in the behaviour of the convergence to the manifold. The constant $\mu_{max} = (2 + K_s)(2 + \chi)$ is chosen to ensure that the growth rate is higher than the death rate at least for some species at the beginning of the dynamics. Such a choice also ensures that all the resources are properly consumed and none of them is growing indefinitely. In all simulations all the other parameters were set to the same values used in the main text.

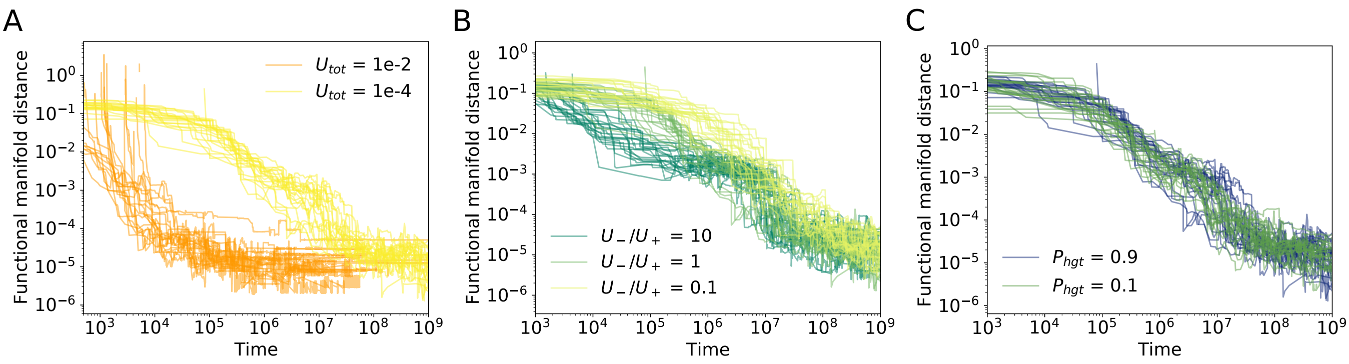


Supplementary Figure S3: The choice of evolutionary parameters does not affect the distance from the functional manifold but can modify the path walked to reach it. 20 realizations for every choice of the parameters are shown. **A** shows the effects of the mutation rate $U_{tot}$. Two values are here considered, a fast mutation rate ($U_{tot} = 10^{-2}$) in orange and a slow one $U_{tot} = 10^{-4}$ in yellow. **B**: effects of the ratio between function loss and function gain rates. Dark green for the case where losing a gene is more probable than gaining it. Light green stands for the even case and yellow for the samples where losing a function is less likely than gaining it. **C**: influence of the probability of gaining new genes via horizontal gene transfer ($P_{hgt}$) versus spontaneous mutation. The distance is calculated as in Fig. S1 In all simulations all the other parameters were set to the same values used in the main text.

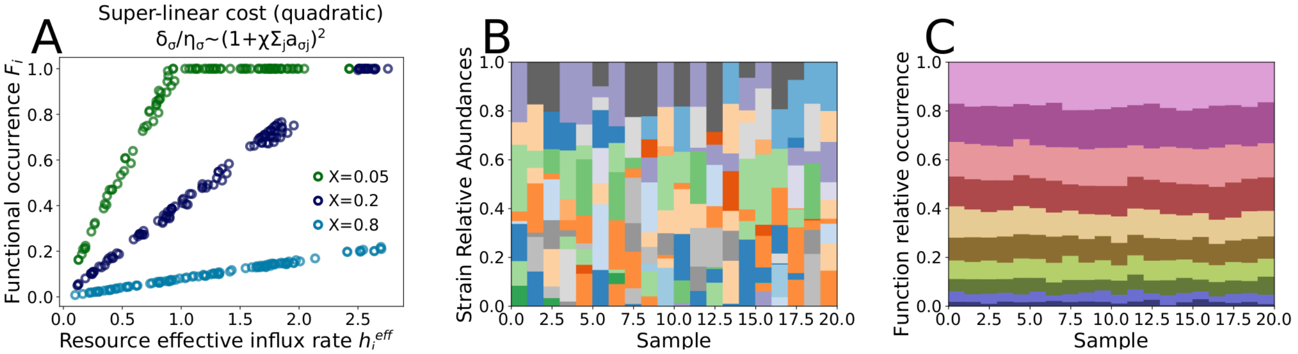


Supplementary Figure S5: Evolutionary outcomes under super-linear metabolic cost. All the panels consider a quadratic cost ($g(z) = (1+z)^2$ in eq. 20). Panel A shows that the functional occurrences $F_i^*$ depend on the effective resource influx rates $h_i^{eff}$ in a similar fashion to what is observed for the linear metabolic cost (see Fig. 2). Different points correspond both to different resources and different realizations of the intrinsic fitness values. Similar to the linear cost, increasing the value of the cost per resource $\chi$ decreases the number of core resources. The other panels show the taxonomic (panel B) and functional (panel C) composition of different communities evolved in independent environments, characterized by the same effective resources influx rate $h_i^{eff}$ but different intrinsic fitness values $\epsilon_\sigma$. Panel B shows that the taxonomic composition varies widely across realizations, while the functional composition is much more stable and minimally affected by intrinsic fitness variation (Panel C). A color in panel B represents a strain, fully characterized by a given functional preference $a_{\sigma\cdot}$. Colors in panel C represent different functions. The overall qualitative picture that emerges confirms the results obtained in the main text for linear metabolic costs. In all simulations, all the other parameters were set to the same values used in the main text. Panel B and C were obtained with $\chi = 0.5$.

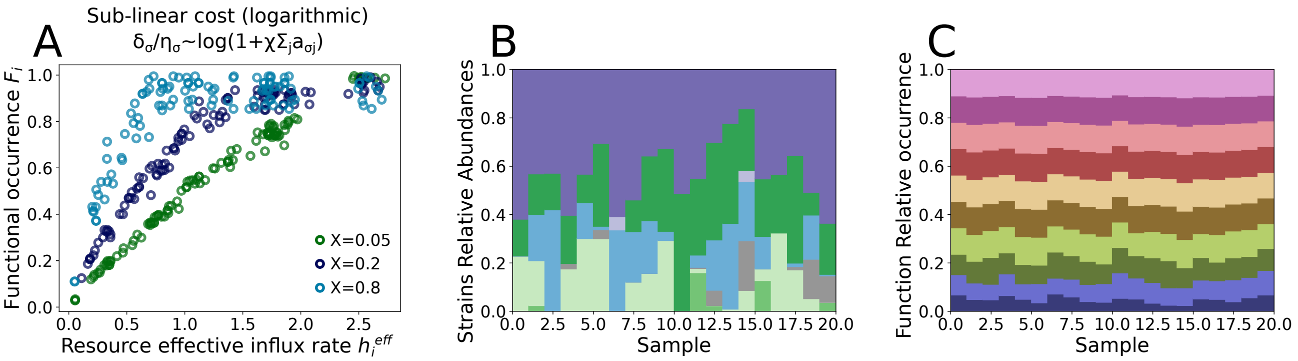


Supplementary Figure S6: Same as figure S5 but with sub-linear metabolic cost. All the panel considers the case of a logarithmic cost ($g(z) = \log(1+z)$ in eq. 20).
